# Supporting Student Learning and Experiences in the Lab: (How) Should We Design Their Groups?

**DOI:** 10.1101/2020.11.19.390658

**Authors:** Tanya Y. Tan, Megan K. Barker

**Affiliations:** Institute for the Study of Teaching & Learning in the Disciplines, Simon Fraser University, Burnaby BC Canada; Department of Biological Sciences, Simon Fraser University, Burnaby BC Canada

**Keywords:** student groups, labs, gender, learning, confidence, skills, introductory biology

## Abstract

Undergraduate science students spend a substantial amount of time working in their laboratory groups, and instructors want to make evidence-based decisions on how to best set up these groups. Despite several studies on group composition, the evidence appears to be quite context-specific, and very little has been published about lab groups. Further, many studies focus solely on conceptual learning; however, the lab is an important venue for also supporting non-content outcomes such as confidence, process skills, team skills, and attitudes. Thus, in our introductory course on molecules, cells, and physiology we were interested in the impact of group composition, on a spread of student outcomes. Students were either placed into groups by the instructor, or self-selected into groups. To assess the impact of group composition on student outcome, we collected pre/post data from >500 students over 2 semesters. Our measures assess conceptual knowledge, confidence in lab skills, attitudes toward group learning, lab grades, gender, year of study, and (via open-ended questions) student perspectives. Using a multiple regression approach, we established models that predict student outcomes based on their individual attributes and on their lab group attributes. Surprisingly, the hetero/homogeneity of the initial group, and whether the groups were student- or instructor-selected, did not affect student outcomes in these models. Further MANCOVA analysis demonstrated that student interaction outside of the lab time was the strongest predictor of positive student attitudes toward group learning. Student perspectives on group formation are mixed, and suggest that a simple and flexible choice approach may best support our students. Overall, these findings have clear implications for our course design and instructional choices: we should focus our efforts to promote positive student interactions, rather than worrying about initial composition.

## Introduction

Two core disciplinary practices identified in undergraduate biology education are collaboration and communication, as described by the AAAS Vision & Change Report [1]. To support our students in developing these skills, groupwork is often used in our courses. Student groupwork has been positively linked to academic achievement, attitudes, and persistence in the discipline (see meta-analysis in [2]). This is consistent with the relatedness component of Self-Determination Theory [3], and with learning as a socially-situated practice [4] – both of which predict positive motivation, growth, and learning as a result of students making peer connections. However, students do not always succeed in cooperative learning; it is critical to structure the groups and group activities in a way that facilitates learning [5–7].

The impact of group composition on student outcomes has been investigated in some contexts: one variable of interest is relevant prior academic achievement or knowledge. This is typically operationalized as pre-requisite grades or a pre-score on a concept inventory. Some studies recommend academically heterogeneous groups -- students grouped together with others who have with a diversity of pre-scores [8,9]. Others recommend groups that are homogeneous in composition [10–13]. Still others note that the academic pre-achievement is not an important factor [14,15].

In addition, demographic attributes are widely used to determine groups, as a practical strategy that does not require collecting/analyzing survey data or incoming quizzes [16,17]. Demographic composition can impact metrics of student learning and behaviours [18–22] – or not [17,23] – with no universal recommendation. The variability of the published recommendations illustrates that the context is quite important when determining optimal group composition. In our search of the literature, we have not found any strong recommendations specifically for undergraduate lab group composition.

In introductory biology course in this study, students work in formal, semester-long lab groups of four. The high-enrollment course includes a diverse population, and for many students it is their first introduction to collaboratively doing science at the university level. In addition to mirroring disciplinary practices, these groups can be important for providing community and supporting first-year students in their transition to university. Historically in this course, these groups are self-selected by students on the first lab day – largely due to convenience and a lack of strong guidance from the literature. With an aim to improve our group-forming process in this course, we systematically investigated the question of group composition on student outcomes. Our goal was to understand and positively impact student performance, confidence, and/or experience in this critical time of transition to university. In this project we investigated the following four research questions:

1. **Impact of initial individual and group scores:** What initial individual and group factors predicted student outcomes? Outcomes were operationalized as a) academic performance, b) content knowledge on a diagnostic test, c) attitude towards groups, and d) confidence in their lab skills.
2. **Demographic differences and who selects the groups**: How does lab group composition differ between student-selected groups and instructor-selected groups? How does a group’s gender composition impact outcomes?
3. **Student interaction outside of lab:** Does outside-lab student-student interaction play a role in student learning? If so, does the group’s gender composition affect students’ tendency to interact outside the lab?
4. **Preferences:** What are student preferences for how groups should be chosen?

## Methods

### Context and Cohort

The participants were students enrolled in a semester-long, high-enrollment introductory biology lab/lecture course at a large university in the Pacific Northwest, over two semesters in 2017. Weekly class contact consisted of three hours of face-to-face lecture, a one-hour tutorial led by a teaching assistant, and a two-hour laboratory supported by the same teaching assistant and a course instructor. Students were invited to participate in the study by completing a biology concept inventory test and survey items that assess self-perceived confidence and attitudes towards group learning at the beginning and end of the academic term. Students who participated in the surveys were given a 0.5% grade bonus. Our dataset comes from 572 total students who consented to be involved in the study, and for whom we obtained both pre- and post-data.

The data used in this article were collected to inform course quality improvement, with consultation from the Simon Fraser University’s Research Ethics Board. Under Article 2.5 of Canada’s Tri-Council Policy Statement on research ethics, quality improvement activities are not subject to institutional ethical review.

### Measures and analysis

A set of measures were used to assess student attributes, including demographics and outcomes, as summarized in Table 1. These individual measures were then used to characterize the group environment that students experienced. For each group, the averages and standard deviations of each of the knowledge test, confidence, and attitude measures were calculated using pre-data. The group average was intended to characterize the group’s benchmark value of the measure while the standard deviation was used to characterize the heterogeneity within the group. Groups were either instructor-selected or student-selected.

**Table 1.** Measures used in this study to capture student attributes, and calculate group attributes. Survey instruments are provided in Supplemental Materials 3.

| **Student Outcome** | **Operational Measure** | **Description / Data Source** |
| --- | --- | --- |
| Lab performance | Lab exam grade | Practical exam, on techniques and material from lab; 12 short-response questions at end of semester. |
| Learning (concept knowledge) | Concept inventory score | 17-item multiple-choice inventory compiled from several relevant sources also used in [24]. Pre/post. |
| Attitude towards group learning | SAGE survey: Student Attitude to Group Environments | 53 Likert-scale questions (from [25]); Pre/post. |
| Confidence in lab skills | Confidence survey | Likert-scale questions; self-rated confidence on lab skills; Pre/post. |
| Demographics | Gender, program level | Gender is a self-reported variable.<br>Program level is from the institute, ranging from 1 to 8; most first-year students were in Level 1 (1-15 credits) or Level 2 (16-30 credits). |
| Familiarity/ interactions with group | Prior familiarity with group (pre);<br>Interaction with group (post) | Both are self-reports. Prior-familiarity was a on a scale of 1 (“never met before” to 3 “knew all group members before”).<br>Interaction with group was a multiple-choice question asking “how often did you work with your lab group?” with the following options: “Labs only, Labs and study time, Labs and lectures, Labs, Lectures and Study time.” |
| Opinion about group formation | Two survey questions (closed- and open-ended) | “In this lab, having instructor-assigned groups is (choose one: better/worse/no different) than having student-selected groups. Please explain your opinion.” Pre/post. |

To determine which initial factors (individual and group) predict student outcomes, we used regression analyses [26,27]. For further analysis of demographics and group relationships, MANCOVA analysis was used [28]. The fuller analysis approach for each of the four study questions is given below in the results section.

### Group Formation

Lab sections were randomly assigned to either the ‘instructor-selected’ or ‘student-selected’ condition. In instructor-selected sections, the instructor assigned groups using student pre-attributes (concept inventory pre-score and gender). As best as possible given the constraint of lab section size and some lack of initial data at the time, the instructor aimed to have an even spread of homogeneous-knowledge (high, mid, and low) and heterogeneous-knowledge groups, without isolating women (based on [21]). In student-selected sections, students formed their own groups on the first day of lab, with no guidance from the instructor aside from the group size. Anecdotally, this tended to be the groups of four people that happened to be sitting together.

## Results

### Descriptive statistics of student measures

Figure 1 presents student scores/responses on knowledge tests, questionnaires and lab exam scores. As a cohort over the semester, students demonstrate increases in their knowledge test scores, confidence in their lab skills, and attitude towards group learning.

**Figure 1.**
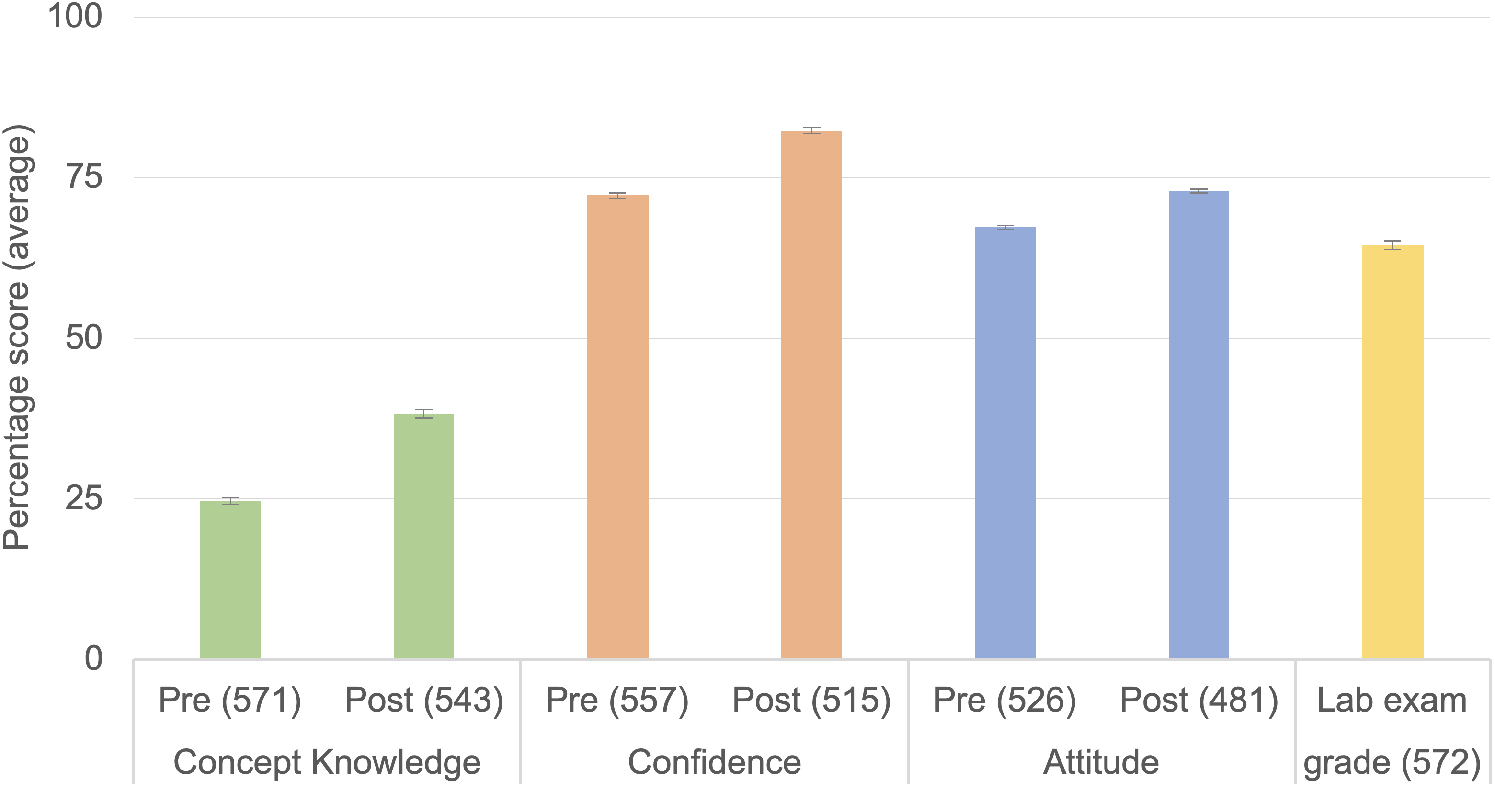
Descriptive statistics of pre- and post-measures. Student scores and responses on measures of knowledge, confidence, attitude toward groups, and the lab exam. Error bars are SEM; Sample sizes (n) given in parentheses. Sample size varies across the measures due to invalid responses to some items or course withdrawals during the term.

### Research Question 1: individual and grouping scores that predict student outcomes

Our goal is to best support student outcomes by thoughtfully designing groups. We thus first needed to determine which initial factors (individual and group) predict student outcomes. We used regression analyses. Supplemental Materials 1 shows the bivariate correlations of the major variables; no collinearity was found between variables. Four multiple regression analyses were performed separately to predict the four end-of-course measures (post-scores): knowledge (concept inventory) scores, attitude (towards group learning), confidence (in lab skills), and lab exam grade. The predictor variables used were the following, entered in two blocks: 1) Individual measures at the start of the course (pre-scores): knowledge, attitude, confidence; and 2) Group attributes at the start of the course (group pre-score average and standard deviation): knowledge, attitude, confidence.

Results from these models are summarized below in Table 2, with the details of each model in Supplemental Materials 2. From these findings, only individual characteristics are significant predictors of student outcomes. Each given pre-variable positively and significantly predicts its own post-variable, which is unsurprising – for example, students with the highest confidence at the start of the course also have the highest confidence at the end. Notably, no group characteristics (average and variability of attributes) predicted any student outcomes.

**Table 2.** Results of multiple regression analysis to determine predictors (X) of student outcomes (Y). In the table, + and - indicate significant (p<0.05) predictors that are positive or negative, respectively. Full model results are shown in Supplemental Materials 2.

| **X =**<br>**Initial variables** |  | *Model 1*<br>**Y =**<br>**Lab exam**<br>**score** | *Model 2*<br>**Y =**<br>**Post-**<br>**knowledge** | *Model 3*<br>**Y =**<br>**Post-**<br>**attitude**<br>**to groups** | *Model 4*<br>**Y =**<br>**Post-**<br>**confidence** |
| --- | --- | --- | --- | --- | --- |
| **Individual...** | **Content knowledge** | + | + |  |  |
|  | **Attitude to groupwork** | + | - | + |  |
|  | **Confidence** |  |  |  | + |
| **Group...**<br>**(hetero- or**<br>**homogeneous)** | **Content knowledge** |  |  |  |  |
|  | **Attitude to groupwork** |  |  |  |  |
|  | **Confidence** |  |  |  |  |

### Research Question 2: Group selection by instructor or students, gender distribution, and impact on outcomes

We explored the difference among female-only, male-only, and mixed gender groups – whether formed by students or by instructors. Overall, the gender distributions across sections are similar (Figure 2). However, when students choose their own groups, the distribution differed: female students tended to form more gender-homogeneous groups than expected from a random sample.

**Figure 2.**
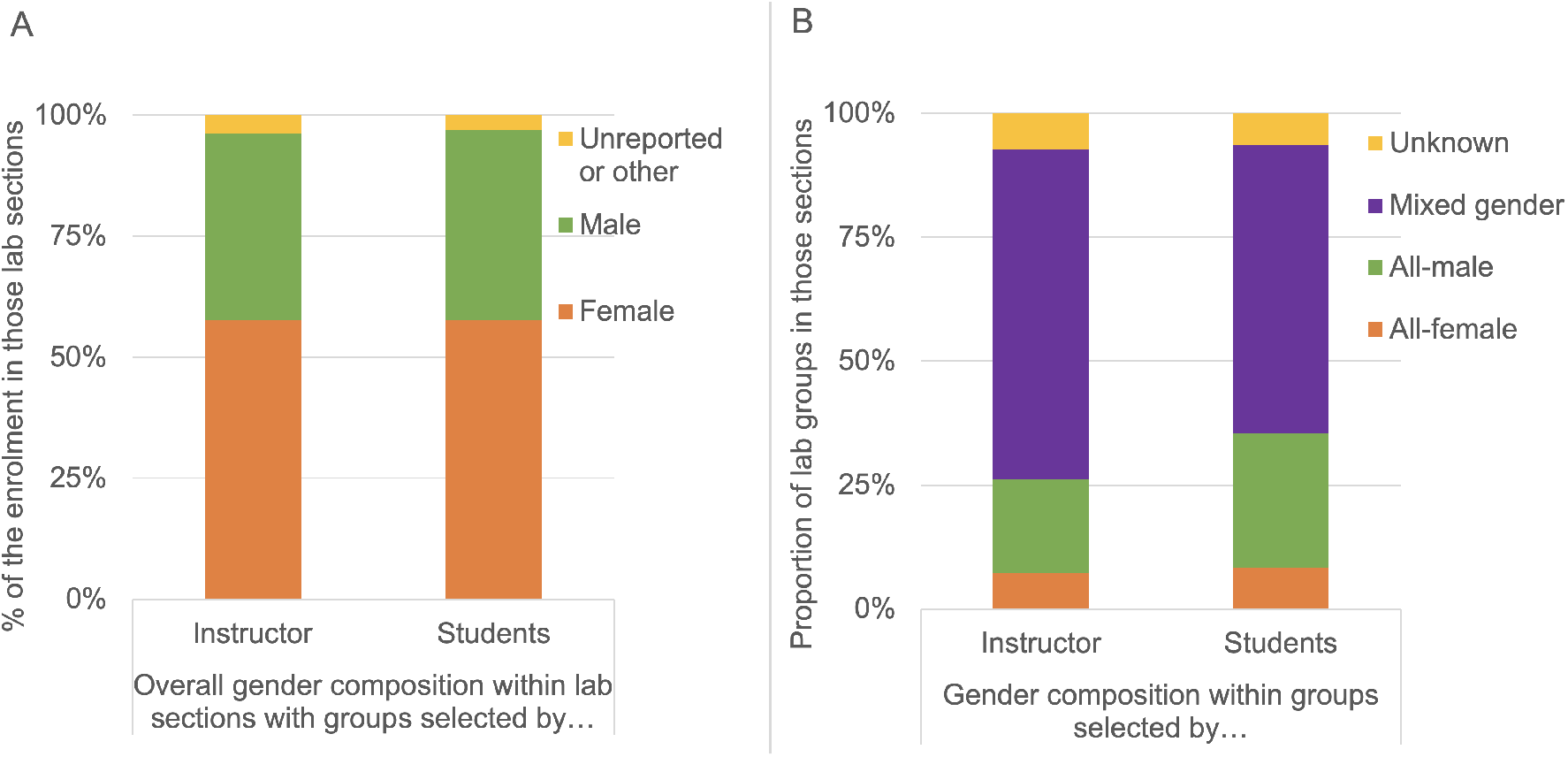
Students self-select into more homogeneous-gender groups than present in instructor-selected groups. *Panel A:* All lab sections had roughly the same gender composition, and so this did not influence group selection type. *Panel B:* Gender composition of lab groups in sections designated for instructor-selected or student-selected groups.

MANCOVA (multivariate analysis of covariance) was conducted to investigate the differences in the learning outcomes of female students on multiple individual post-measures (knowledge, attitude, confidence, lab exam grade) by the gender composition (either in female only or mixed-gender groups). The individual pre-measures are controlled as covariates, including knowledge, attitude, confidence scores. The impact of female only vs. Mixed-gender groups, is however, not significant *(F(4, 180) = 1.138, p = 0.340, Wilk’sΛ = 0.975, partial η^2^ = 0.025).* We conducted the same analysis on male students, comparing their performance in male-only and mixed-gender group, and found that the gender group composition doesn’t have a statistically significant impact on their learning outcome either (*F*(4, 113) = 1.110, p = 0.355, Wilk’s Λ = 0.962, partial η^2^ = 0.038).

### Question 3: Impact of group interaction on outcomes; impact of gender on student interaction

To understand one aspect of group dynamics, students were asked how often they interacted with the group, and how well they knew the group (n =420). 88.3% of the students “had never really met or worked with the other group members before,” 9.8% “knew some of the other group members,” and only 1.9% knew all the group members. Despite that most students did not know their group members at the beginning, 55.0% of the students interacted with their group outside the laboratory. The patterns of interaction varied between the two semesters studied, and from prior instructor experience, summer enrollment has quite a different population (returning students and early keeners) compared to fall (first semester at university). The fall semester also has the highest semesterly enrollment for this course, so the rest of the interactions were analyzed using only this data.

We examined the impact of outside-lab-interaction with student outcomes. We used MANCOVA to assess statistical differences on multiple post measures (knowledge, confidence attitude,) by an independent grouping variable: whether or not students interacted with their group members outside the lab, and how well students knew their group, separately. All individual pre-measures were used as the covariates: knowledge, confidence, attitude, that the analysis controlled for.

There was a statistically significant difference in the outcome variables based on whether one interacted with the group outside the lab, *F (3, 379) = 9.045, p < .001; Wilk’sΛ = 0.933, partial η2 = 0.067.* The between-subjects tests confirm that, interactions outside the lab have a significant impact (adjusted p<0.05) on post-attitude but not on post-knowledge or post-confidence scores. Note here we used alpha = 0.05/3 as the threshold value following the Bonferroni correction.

A second MANCOVA was performed to examine the difference on multiple group-level post measures (group means of post-knowledge, -attitude, -confidence; and lab exam scores) by the grouping variable: whether or not the group has at least half of the students reporting interacting outside the lab. The group-level pre measures (means of confidence, attitude, knowledge) were used as covariates to control for any difference among the groups at the beginning of the term. In this analysis, 47 (40.2%) groups were identified as non-interactive while 70 groups (59.8%) were identified as interactive. A significant difference was found between the interactive and noninteractive group: *F(4, 109) = 3.891, p = 0.005, Wilk’s Λ = 0.875, partial η^2^ = 0.125.* The between-subjects tests confirmed that whether the group is interactive has a significant impact on the group’s average Post Attitude, *F(1,112)= 14.112, p<0.001, partial η^2^ = 0.112.* However, there is no significant difference on other outcome variables (knowledge, confidence, lab exam scores; p>0.05).

We then investigated the what promoted outside-lab student interaction. In 1177 cohort, since 83.6% of the students “had never really met or worked with the other group members before”, the initial familiarity with group members isn’t a determining factor of outside lab interaction. A Chi-square test of independence is performed to determine whether there is an association between categorical variables: outside lab interaction, hetero/homogeneous groups in terms of level and gender. The associations were marginally significant (non-significant) between the outside lab interaction and gender composition (χ2 (2)=5.754, p=0.056 while the association between outside lab interaction and level composition (χ2 (2)=1.140, p=0.434) was not statistically significant. For female students, 58.5% of the female students in the female-only group had outside lab interaction while only 50.7% of the female students in mixed-gender group had outside-lab interaction. Similarly, for male students, 74.1% of the students in male-only group had outside lab interaction, while in mixed gender group, only 53.6 % of the male students had outside lab interaction with their peers. This might suggest students’ increased tendency to have outside lab interaction when they are in a single gender group. Individual’s gender and level have no detectable association with their outside lab interaction (p =0.236 and 0.694 respectively from Chi-squared tests). Nor does the instructor-selected/student selected have an effect (p=0.427), even though students tended to form gender-homogeneous groups slightly more often (9% more single-gender groups than in instructor-selected sections).

Further, a logistic regression was performed to ascertain the effects of pre-attitude and group gender composition on the likelihood that participants would have outside lab interaction. The model is statistically significant, χ2 (3)= 23.737, *p* < 0.001, Nagelkerke R^2^ = 0.086, indicating that the model explains 8.6% of the variance in the dependent variable (outside lab interaction). Both Pre-Attitude (Wald = 16.092, df=1, p<0.001) and Group’s gender composition (Wald = 6.847, df=2, p=0.033) were statistically significant predictors. The full model for the probability (p) of outside lab interaction is: ln(p/(1-p)) = −4.043 + 0.023 Pre-Attitude Total + 0.429 x_1_+1.196x_2_, where x_1_=1 for female only groups, x_2_=1 for male only groups, and mixed-gender groups have x_1_=0 and, x_2_=0 The model was able to predict 62% of the cases compared to 56% in the constant only model.

Gender impact on interactions is graphed in Figure 3: both homogeneous-gender groups were more likely to have outside lab interactions than mixed-gender groups, controlling for Pre-Attitude. The odds ratio for pre-attitude total score is 1.023, indicating for one unit of increase in Pre-Attitude scores, it increases the odds of outside lab interaction by 2.3%. This is statistically significant, but not a large impact.

**Figure 3.**
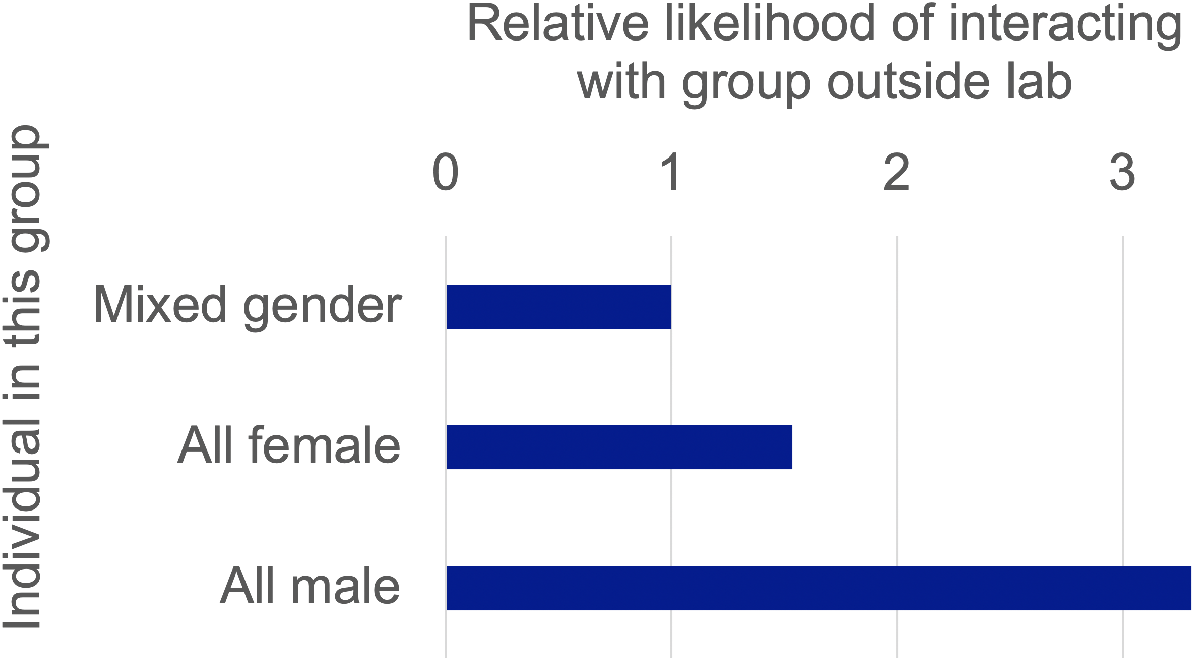
Homogeneous-gender groups interact outside of lab more than mixed-gender groups. Results from logistic regression; values shown are odds ratios.

In summarizing the findings from research questions 2 and 3, there are two main points to consolidate. First, groups that interacted outside the lab were more likely to have an end-of-course positive attitude toward group learning. This interaction did not, however, impact content learning, lab exam performance, or confidence in lab skills. Secondly, gender-homogeneous groups were more likely to interact outside the lab.

### Question 4: Student preferences for group selection by students or instructors

On the pre- and post-survey, students were asked who should choose the lab groups (students, instructors, or no preference), and given space to explain their reasoning. These data are presented visually (Figure 4), using matched students who completed both the pre and the post question. This allows us to see a general view of which students change opinions, in which conditions. The open-response question was summarized for general themes, looking for commonalities in student responses.

**Figure 4.**
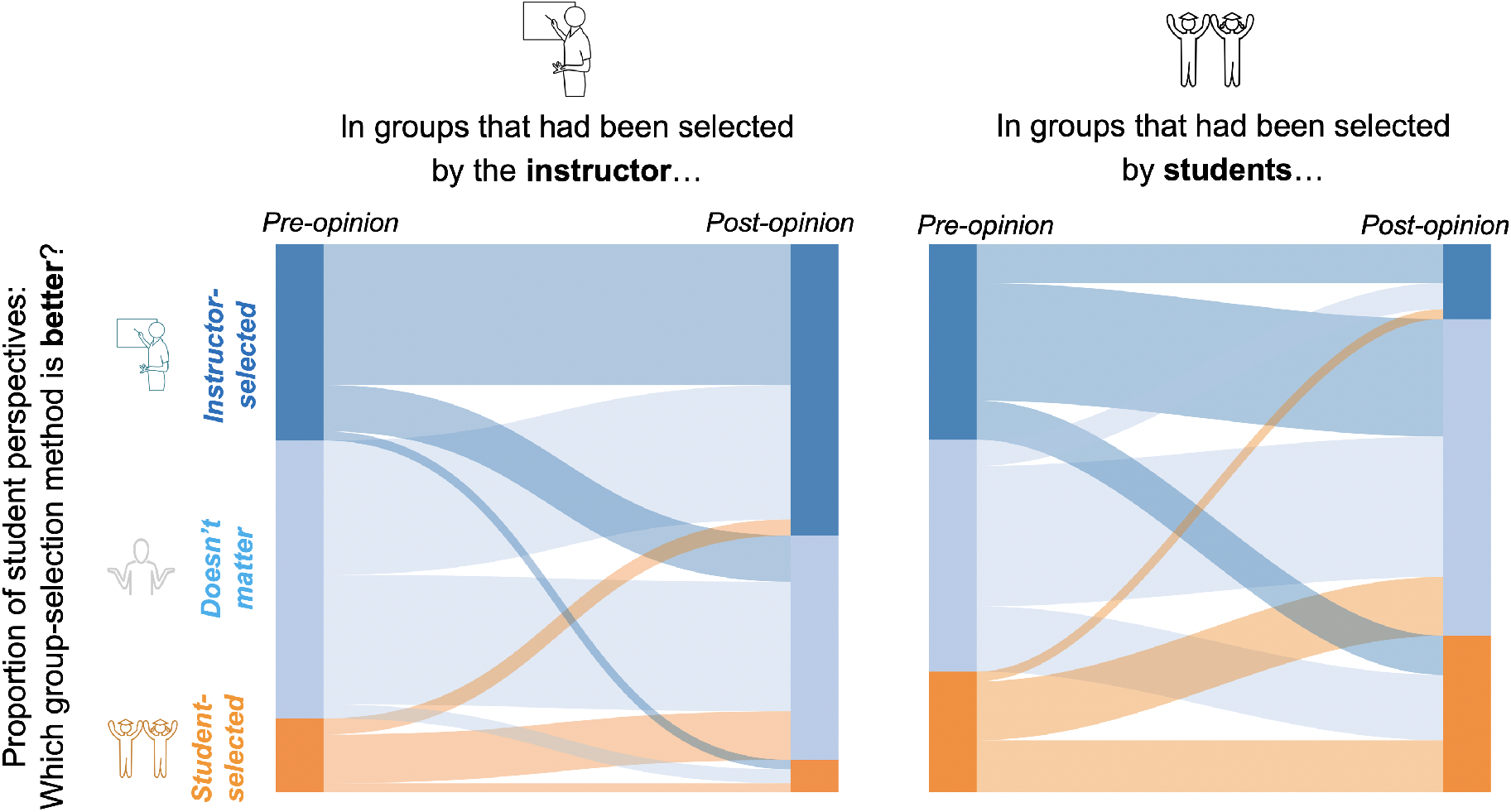
Student perspectives of who *should* choose the group may relate to who *did* choose their group. Sankey diagram representing proportions of the class with given perspectives at start and end of term, in either of the two setups (student-selected or instructor-selected group formation). Flow shows opinions that changed or remained consistent. Data includes responses from students who answered the question both at the start and end of the semester; n=237 students from instructor-selected groups and n=168 students from student-selected groups.

From a visual analysis of Figure 4, both populations are reasonably similar at the start of term, with somewhat more student-selection-preference in the student-self-selection lab sections. Very few students switched from one strong opinion to another. When students have chosen their own groups, a large proportion of them report that it didn’t matter who had chosen the groups; this differs from the instructor-determined groups. Students who didn’t care at the start of term had different opinions at the end; many more in the instructor-decided sections would have preferred for the instructor to choose.

Students reported the reasoning behind their perspectives as to who should determine groups, with a spread of perspectives. From an overview of student responses, several comments were consistent, and are reported in Table 3.

**Table 3.** Summary of students’ reasoning for their perspectives on group choice.

| **Who should choose the groups?** | **Why?** |
| --- | --- |
| The instructor | Didn’t know anyone; want to meet new people; less distracting to not work with friends; takes social pressure off students. |
| It doesn’t matter | Didn’t know anyone at start; it’s up to each group, so it can go well or badly either way. |
| The students | Want freedom to choose; more comfortable; know how to work/communicate with friends. |

## Discussion and Conclusions

### From an initial question: Who should choose the groups?

An initial project question centred around the differences in student outcomes from instructor-selected vs. student-selected group formation. However, course enrollment changes and section switches essentially made this operationally/logistically moot. Further, as the overwhelming number of the students reported that they had never met their groups before, and as this was a first-semester, first-year course, it did not appear to be relevant. Thus, our analysis followed an ‘action research’ approach, and focused instead on the outcomes of groups based on their initial composition, and thus whether or not to use initial student variables to choose the groups at all.

### What group composition has the best student outcomes?

We used regression data to predict student outcomes based on individual and group attributes. The initial individual scores-based variables (attitude to groups, content knowledge, confidence in lab skills) each predicted their own post-scores. These individual variables are not under the control of the instructor. The group scores-based variables (average and variability on attitude, content knowledge, confidence) did not predict any student outcomes. This was surprising, and it suggests that these group characteristics, in bulk, do not impact a student’s experience in our lab course. Thus, from an instructional perspective, in our context, it is not worth designing groups using these metrics.

Logistically speaking, in a course where there are no shared grades among group members, there is little motivation for students to work strongly with their group unless they already have positive interactions together. As discussed before, the context is a hugely important factor in student and group outcomes. Thus, these findings are likely not generalizable to other courses – such as where the groupwork is a direct, graded course component [29]; or at different academic year levels.

### Group selection: a responsibility or an opportunity?

From a visual analysis of our students’ perspectives, most seemed to prefer whichever group-formation method they experienced, or they do not have a strong opinion. In this context, it is unsurprising – this is a first-year, first-semester course. There was reasonable reasoning in support of both group-formation ‘ownership’ (student or instructor). One student commented that “the instructor should choose, because then there’s no chance [for the student] to blame oneself for choosing the wrong group this way.” This philosophy could be applied in either direction – an instructor may want to avoid strong complaints and the perceived arrow of blame for how the group interaction fares. Anecdotally, when the instructor chose the groups, the instructor noted receiving a non-trivial number of emails with strong negative opinions. Students may have been having similar experiences when they selected their own groups, but they were not reaching out to the instructor for support or solutions. In this way, the group-selection process may be an indicator of instructor-student immediacy [30]. So, the choice also depends on the instructional goal-avoiding unresolvable complaints and encouraging self-resolution of group issues, or being connected to and actively supporting the student experience in one’s course.

### Student interaction impacts attitude, and gender may impact interaction

Student self-reporting that they interacted with their group was a positive predictor of their attitude to groupwork (but not other outcome measures). This analysis controlled for other initial variables, and so we can interpret that group interaction is a benefit to these students. We also note that, in our dataset, gender composition of the group predicts interaction. A preliminary instructional response would then be to support (or design) gender-homogeneous groups. However, this contrasts with the current, important pedagogical trends towards supporting diversity and integration, rather than segregation, in our classrooms. Further, gender grouping has community implications as well: gender has impacted the inaccurate perception of intelligence [31] and social status [32] in biology classes. As described in the Vision & Change Report for Undergraduate Biology [1], our students “need to develop skills to participate in diverse working communities, as well as the ability to take full advantage of their collaborators’ multiple perspectives and skills.” Thus, we would be doing our students a broader disservice by segregating by gender. This equity is important, as is our essential instructional role in supporting students to be comfortable in these heterogeneous groupings [33,34].

### A goal shift: promoting student interaction and community

In practice, what to do with all of this? To address student anxiety about group formation, and reduce logistical concerns, we could simply randomize groups on day one, and allow for flexibility when enrollment changes. Based on the findings here, our instructional team has chosen a hybrid approach that is also consistent with principles of Universal Design for Learning (such as allowing choice) [35]. We give students advance notice that they’ll have the option to choose their own groups if they so wish; and for any who prefer not to (or don’t declare), we will help them find a group. Further study will need to evaluate the effectiveness of this approach.

One additional instructional comment is that non-random group assignment for a course this size was not a trivial task. First of all, it needed to be done quickly (collecting and compiling student attributes in the first week of the course). Second of all, the number of students who dropped or added the course, or switched lab sections, substantially disrupted the process. As a result, it is almost a relief that the evidence-based instructional choice is to not worry about initial group composition in our context.

More meaningfully, rather than focus efforts on the initial group composition, in this course our instructional role is to find ways to support group interaction in a meaningful and useful way. Some ideas from the literature on Team-Based Learning, Co-operative Learning, and group development processes [7,36–38]would be useful to structure these course supports. Initial ideas include structuring group process; building dedicated course time for groups to move through group development stages; and increasing the difficulty of shared tasks to require positive group interdependence. Regardless of the group that a given student ends up in, the more we can support their belonging, the more we can positively impact their motivation, growth, and learning.

## Acknowledgments

The authors gratefully acknowledge Onkar Bains and Peter Hollmann, course co-instructors, for their support in survey distribution and discussion. We also are grateful to the Simon Fraser University Institute for the Study of Teaching & Learning in the Disciplines, including Laura D’Amico, Sheri Fabian, and Tara McFarlane, for helpful advice on experimental design and for research assistantship support.

