## Supplemental Materials for "Supporting Student Learning and Experiences in the Lab: (How) Should We Design Their Groups?"

### Supplemental Materials 1: Correlations

The table below shows the bivariate correlations of the major variables. No collinearity was found between variables. It should be noted that students' knowledge test scores are weakly correlated with their confidence scores at the beginning and end of the term which indicates students' low levels of metacognitive ability of one's learning in the first year. The knowledge test scores also weakly correlated with attitude measures ($r<0.2$). Overall, the pre measures are moderately correlated with the corresponding post-measures which means, the academic ability, confident and attitude of the students at the start of the term has an impact on these traits at the end of the term, but only moderately. (pairwise exclusion)

**Correlations**

| | | Lab Exam Percent Score | Post Attitude Total | Post Confidence Total | PostCI score | Pre Attitude Total | Pre Confidence Total | PreCIscore |
|---|---|---|---|---|---|---|---|---|
| Lab Exam Percent Score | Pearson Correlation | 1 | -.065 | .142** | .471** | -.168** | .004 | .222** |
| | Sig. (2-tailed) | | .154 | .001 | .000 | .000 | .929 | .000 |
| | N | 572 | 480 | 514 | 542 | 510 | 542 | 555 |
| Post Attitude Total | Pearson Correlation | -.065 | 1 | .123** | -.045 | .543** | -.041 | -.115* |
| | Sig. (2-tailed) | .154 | | .008 | .340 | .000 | .374 | .013 |
| | N | 480 | 481 | 466 | 455 | 441 | 463 | 470 |
| Post Confidence Total | Pearson Correlation | .142** | .123** | 1 | .025 | .075 | .530** | .051 |
| | Sig. (2-tailed) | .001 | .008 | | .579 | .106 | .000 | .255 |
| | N | 514 | 466 | 515 | 488 | 466 | 496 | 503 |
| PostCI score | Pearson Correlation | .471** | -.045 | .025 | 1 | -.140** | .001 | .351** |
| | Sig. (2-tailed) | .000 | .340 | .579 | | .002 | .978 | .000 |
| | N | 542 | 455 | 488 | 543 | 484 | 515 | 533 |
| Pre Attitude Total | Pearson Correlation | -.168** | .543** | .075 | -.140** | 1 | .060 | -.106* |
| | Sig. (2-tailed) | .000 | .000 | .106 | .002 | | .177 | .017 |
| | N | 510 | 441 | 466 | 484 | 526 | 517 | 512 |
| Pre Confidence Total | Pearson Correlation | .004 | -.041 | .530** | .001 | .060 | 1 | .100* |
| | Sig. (2-tailed) | .929 | .374 | .000 | .978 | .177 | | .020 |
| | N | 542 | 463 | 496 | 515 | 517 | 557 | 542 |
| PreCIscore | Pearson Correlation | .222** | -.115* | .051 | .351** | -.106* | .100* | 1 |
| | Sig. (2-tailed) | .000 | .013 | .255 | .000 | .017 | .020 | |
| | N | 555 | 470 | 503 | 533 | 512 | 542 | 571 |

**. Correlation is significant at the 0.01 level (2-tailed).

*. Correlation is significant at the 0.05 level (2-tailed).

- Pre and its matching post measures (e.g. PreCI and postCI) generally correlate
- PreAttitude is negatively correlated with preCI however, the correlations between Post-Attitude and PostCI, /Lab Exam is not significant


Attribution CC-BY-NC-ND.

### Supplemental Materials 2: Regression Models

#### *2.1 Lab Exam Scores: Only pre-CI and pre-Attitude are significant predictors*

**Model Summary[d]**

| Model | R | R Square | Adjusted R Square | Std. Error of the Estimate | Change Statistics: R Square Change | F Change | df1 | df2 | Sig. F Change | Durbin-Watson |
|---|---|---|---|---|---|---|---|---|---|---|
| 1 | .264[a] | .070 | .064 | 15.07365623 | .070 | 11.924 | 3 | 478 | .000 | |
| 2 | .281[b] | .079 | .067 | 15.04460737 | .009 | 1.616 | 3 | 475 | .185 | |
| 3 | .290[c] | .084 | .067 | 15.05069930 | .005 | .872 | 3 | 472 | .456 | 1.683 |

a. Predictors: (Constant), preConfidenceTotal, preAttitudeTotal, PreCIscore

b. Predictors: (Constant), preConfidenceTotal, preAttitudeTotal, PreCIscore, preConfidenceTotal_groupavg, preAttitudeTotal_group_avg, PreCIscore_groupavg

c. Predictors: (Constant), preConfidenceTotal, preAttitudeTotal, PreCIscore, preConfidenceTotal_groupavg, preAttitudeTotal_group_avg, PreCIscore_groupavg, PreConf_GroupSD, preAttitudeTotalgroupSD, PreCIscoregroupSD

d. Dependent Variable: LabExamPercentScore

**Coefficients[a]**

| Model | | Unstandardized Coefficients B | Std. Error | Standardized Coefficients Beta | t | Sig. |
|---|---|---|---|---|---|---|
| 1 | (Constant) | 79.458 | 7.665 | | 10.367 | .000 |
| | preAttitudeTotal | -.125 | .034 | -.161 | -3.616 | .000 |
| | PreCIscore | 25.175 | 5.830 | .193 | 4.318 | .000 |
| | preConfidenceTotal | .020 | .092 | .010 | .221 | .825 |
| 2 | (Constant) | 106.267 | 14.427 | | 7.366 | .000 |
| | preAttitudeTotal | -.088 | .041 | -.113 | -2.128 | .034 |
| | PreCIscore | 28.037 | 8.815 | .215 | 3.181 | .002 |
| | preConfidenceTotal | .085 | .108 | .041 | .791 | .429 |
| | preAttitudeTotal_group_avg | -.130 | .076 | -.091 | -1.716 | .087 |
| | PreCIscore_groupavg | -6.577 | 11.980 | -.037 | -.549 | .583 |
| | preConfidenceTotal_groupavg | -.250 | .202 | -.064 | -1.235 | .218 |
| 3 | (Constant) | 110.764 | 14.867 | | 7.450 | .000 |
| | preAttitudeTotal | -.089 | .041 | -.115 | -2.168 | .031 |
| | PreCIscore | 27.844 | 8.823 | .213 | 3.156 | .002 |
| | preConfidenceTotal | .084 | .108 | .040 | .776 | .438 |
| | preAttitudeTotal_group_avg | -.142 | .076 | -.100 | -1.861 | .063 |
| | PreCIscore_groupavg | -8.274 | 12.187 | -.047 | -.679 | .498 |
| | preConfidenceTotal_groupavg | -.259 | .203 | -.066 | -1.274 | .203 |
| | PreConf_GroupSD | -.120 | .233 | -.023 | -.514 | .608 |
| | preAttitudeTotalgroupSD | -.087 | .079 | -.050 | -1.099 | .272 |
| | PreCIscoregroupSD | 17.063 | 14.084 | .057 | 1.212 | .226 |

a. Dependent Variable: LabExamPercentScore


Attribution CC-BY-NC-ND.

### *2.2 Content knowledge ("CI") Scores: Only pre CI is a significant predictor*

**Model Summary**[d]

| Model | R | R Square | Adjusted R Square | Std. Error of the Estimate | Change Statistics | | | | | Durbin-Watson |
|---|---|---|---|---|---|---|---|---|---|---|
| | | | | | R Square Change | F Change | df1 | df2 | Sig. F Change | |
| 1 | .368[a] | .135 | .130 | .154376373 | .135 | 26.299 | 3 | 504 | .000 | |
| 2 | .378[b] | .143 | .133 | .154162827 | .008 | 1.466 | 3 | 501 | .223 | |
| 3 | .383[c] | .147 | .131 | .154285126 | .004 | .735 | 3 | 498 | .531 | 1.821 |

a. Predictors: (Constant), preConfidenceTotal, PreCIscore, preAttitudeTotal

b. Predictors: (Constant), preConfidenceTotal, PreCIscore, preAttitudeTotal, preConfidenceTotal_groupavg, preAttitudeTotal_group_avg, PreCIscore_groupavg

c. Predictors: (Constant), preConfidenceTotal, PreCIscore, preAttitudeTotal, preConfidenceTotal_groupavg, preAttitudeTotal_group_avg, PreCIscore_groupavg, PreConf_GroupSD, preAttitudeTotalgroupSD, PreCIscoregroupSD

d. Dependent Variable: PostCIscore

**Coefficients**[a]

| Model | | Unstandardized Coefficients | | Standardized Coefficients | t | Sig. |
|---|---|---|---|---|---|---|
| | | B | Std. Error | Beta | | |
| 1 | (Constant) | .307 | .073 | | 4.220 | .000 |
| | preAttitudeTotal | .000 | .000 | -.036 | -.867 | .386 |
| | PreCIscore | .493 | .057 | .363 | 8.698 | .000 |
| | preConfidenceTotal | .000 | .001 | .012 | .283 | .778 |
| 2 | (Constant) | .371 | .125 | | 2.960 | .003 |
| | preAttitudeTotal | .000 | .000 | -.047 | -.911 | .363 |
| | PreCIscore | .381 | .086 | .281 | 4.441 | .000 |
| | preConfidenceTotal | .001 | .001 | .050 | 1.007 | .314 |
| | preAttitudeTotal_group_avg | .000 | .001 | .012 | .239 | .811 |
| | PreCIscore_groupavg | .195 | .113 | .108 | 1.718 | .086 |
| | preConfidenceTotal_groupavg | -.003 | .002 | -.069 | -1.396 | .163 |
| 3 | (Constant) | .377 | .128 | | 2.941 | .003 |
| | preAttitudeTotal | .000 | .000 | -.050 | -.952 | .341 |
| | PreCIscore | .374 | .086 | .275 | 4.347 | .000 |
| | preConfidenceTotal | .001 | .001 | .047 | .951 | .342 |
| | preAttitudeTotal_group_avg | 9.476E-5 | .001 | .007 | .140 | .889 |
| | PreCIscore_groupavg | .165 | .117 | .092 | 1.419 | .157 |
| | preConfidenceTotal_groupavg | -.003 | .002 | -.067 | -1.350 | .178 |
| | PreConf_GroupSD | .001 | .002 | .022 | .522 | .602 |
| | preAttitudeTotalgroupSD | .000 | .001 | -.021 | -.498 | .619 |
| | PreCIscoregroupSD | .193 | .142 | .061 | 1.354 | .176 |

a. Dependent Variable: PostCIscore

### *2.3 Attitude towards Group Learning: Only pre-Attitude is a significant predictor*

**Model Summary[d]**

| Model | R | R Square | Adjusted R Square | Std. Error of the Estimate | Change Statistics | | | | | Durbin-Watson |
|---|---|---|---|---|---|---|---|---|---|---|
| | | | | | R Square Change | F Change | df1 | df2 | Sig. F Change | |
| 1 | .572[a] | .328 | .323 | 18.351 | .328 | 72.739 | 3 | 448 | .000 | |
| 2 | .577[b] | .333 | .324 | 18.342 | .005 | 1.146 | 3 | 445 | .330 | |
| 3 | .577[c] | .333 | .319 | 18.400 | .000 | .069 | 3 | 442 | .976 | 1.922 |

a. Predictors: (Constant), preConfidenceTotal, preAttitudeTotal, PreCIscore

b. Predictors: (Constant), preConfidenceTotal, preAttitudeTotal, PreCIscore, preConfidenceTotal_groupavg, preAttitudeTotal_group_avg, PreCIscore_groupavg

c. Predictors: (Constant), preConfidenceTotal, preAttitudeTotal, PreCIscore, preConfidenceTotal_groupavg, preAttitudeTotal_group_avg, PreCIscore_groupavg, PreConf_GroupSD, preAttitudeTotalgroupSD, PreCIscoregroupSD

d. Dependent Variable: postAttitudeTotal

**Coefficients[a]**

| Model | | Unstandardized Coefficients | | Standardized Coefficients | t | Sig. |
|---|---|---|---|---|---|---|
| | | B | Std. Error | Beta | | |
| 1 | (Constant) | 99.035 | 9.119 | | 10.860 | .000 |
| | preAttitudeTotal | .590 | .040 | .569 | 14.610 | .000 |
| | PreCIscore | -8.306 | 6.966 | -.046 | -1.192 | .234 |
| | preConfidenceTotal | -.161 | .113 | -.055 | -1.423 | .156 |
| 2 | (Constant) | 86.304 | 16.002 | | 5.393 | .000 |
| | preAttitudeTotal | .602 | .052 | .580 | 11.601 | .000 |
| | PreCIscore | -12.423 | 10.555 | -.069 | -1.177 | .240 |
| | preConfidenceTotal | -.288 | .137 | -.099 | -2.101 | .036 |
| | preAttitudeTotal_group_avg | -.027 | .088 | -.015 | -.305 | .761 |
| | PreCIscore_groupavg | 8.309 | 13.800 | .035 | .602 | .547 |
| | preConfidenceTotal_groupavg | .412 | .249 | .078 | 1.657 | .098 |
| 3 | (Constant) | 85.633 | 16.413 | | 5.217 | .000 |
| | preAttitudeTotal | .602 | .052 | .580 | 11.552 | .000 |
| | PreCIscore | -12.317 | 10.608 | -.069 | -1.161 | .246 |
| | preConfidenceTotal | -.287 | .138 | -.099 | -2.080 | .038 |
| | preAttitudeTotal_group_avg | -.024 | .089 | -.013 | -.266 | .790 |
| | PreCIscore_groupavg | 8.342 | 14.209 | .036 | .587 | .557 |
| | preConfidenceTotal_groupavg | .403 | .251 | .077 | 1.607 | .109 |
| | PreConf_GroupSD | -.014 | .291 | -.002 | -.047 | .962 |
| | preAttitudeTotalgroupSD | .042 | .097 | .017 | .437 | .663 |
| | PreCIscoregroupSD | -2.463 | 17.668 | -.006 | -.139 | .889 |

a. Dependent Variable: postAttitudeTotal


Attribution CC-BY-NC-ND.

### 2.4 Confidence in Lab Skills: Only pre-confidence is a significant predictor

**Model Summary[d]**

| Model | R | R Square | Adjusted R Square | Std. Error of the Estimate | Change Statistics | | | | | Durbin-Watson |
|---|---|---|---|---|---|---|---|---|---|---|
| | | | | | R Square Change | F Change | df1 | df2 | Sig. F Change | |
| 1 | .503[a] | .253 | .249 | 6.087 | .253 | 53.631 | 3 | 474 | .000 | |
| 2 | .508[b] | .258 | .249 | 6.086 | .005 | 1.068 | 3 | 471 | .362 | |
| 3 | .511[c] | .261 | .247 | 6.095 | .002 | .523 | 3 | 468 | .667 | 1.944 |

a. Predictors: (Constant), preConfidenceTotal, PreClscore, preAttitudeTotal

b. Predictors: (Constant), preConfidenceTotal, PreClscore, preAttitudeTotal, preConfidenceTotal_groupavg, preAttitudeTotal_group_avg, PreClscore_groupavg

c. Predictors: (Constant), preConfidenceTotal, PreClscore, preAttitudeTotal, preConfidenceTotal_groupavg, preAttitudeTotal_group_avg, PreClscore_groupavg, PreConf_GroupSD, preAttitudeTotalgroupSD, PreClscoregroupSD

d. Dependent Variable: postConfidenceTotal

**Coefficients[a]**

| Model | | Unstandardized Coefficients | | Standardized Coefficients | t | Sig. |
|---|---|---|---|---|---|---|
| | | B | Std. Error | Beta | | |
| 1 | (Constant) | 31.720 | 2.987 | | 10.619 | .000 |
| | preAttitudeTotal | .013 | .013 | .039 | .976 | .330 |
| | PreClscore | 1.652 | 2.237 | .029 | .738 | .461 |
| | preConfidenceTotal | .458 | .037 | .496 | 12.449 | .000 |
| 2 | (Constant) | 34.925 | 5.143 | | 6.790 | .000 |
| | preAttitudeTotal | .030 | .017 | .089 | 1.772 | .077 |
| | PreClscore | 1.028 | 3.402 | .018 | .302 | .763 |
| | preConfidenceTotal | .445 | .044 | .482 | 10.139 | .000 |
| | preAttitudeTotal_group_avg | -.046 | .028 | -.082 | -1.646 | .100 |
| | PreClscore_groupavg | 1.246 | 4.524 | .017 | .275 | .783 |
| | preConfidenceTotal_groupavg | .051 | .080 | .030 | .638 | .524 |
| 3 | (Constant) | 34.862 | 5.288 | | 6.593 | .000 |
| | preAttitudeTotal | .029 | .017 | .088 | 1.756 | .080 |
| | PreClscore | .972 | 3.412 | .017 | .285 | .776 |
| | preConfidenceTotal | .445 | .044 | .482 | 10.121 | .000 |
| | preAttitudeTotal_group_avg | -.046 | .028 | -.083 | -1.645 | .101 |
| | PreClscore_groupavg | .316 | 4.646 | .004 | .068 | .946 |
| | preConfidenceTotal_groupavg | .043 | .080 | .026 | .535 | .593 |
| | PreConf_GroupSD | -.007 | .094 | -.003 | -.072 | .942 |
| | preAttitudeTotalgroupSD | .031 | .032 | .040 | .972 | .331 |
| | PreClscoregroupSD | 3.675 | 5.752 | .028 | .639 | .523 |

a. Dependent Variable: postConfidenceTotal


Attribution CC-BY-NC-ND.

### Supplemental Materials 3: Survey Instruments

#### 3.1 Confidence Questionnaire

This part asks you about your confidence in your lab skills, at this stage of our course.
You are asked to rate your confidence in each lab activity by selecting a number between 1 and 5 where the numbers mean the following:

**1= Not at all confident**
2 = Not very confident
3= Neutral
4= Somewhat Confident
**5 = Fully confident**

1. Choose the appropriate lab equipment for a given task or scientific study
2. Prepare a wet-mounted slide, view at appropriate magnification with a compound microscope, and calculate the total magnification
3. Explain the purpose of experimental controls in a given lab procedure
4. Troubleshoot problems that have arisen in an experiment you have done
5. State the independent and dependent variables, given a hypothesis
6. Design an experiment to test a simple hypothesis
7. Summarize raw data in a properly-labeled graph, and interpret that graph using background understanding and common sense
8. Use pipettes of the appropriate size with accuracy and efficiency
9. Analyze numerical data from an experiment you have done, and describe the conclusions
10. Identify anatomical structures on a rat dissection
11. Make a reasonable prediction about the results of a procedure you are about to perform
12. Record detailed observations and notes while performing scientific studies
13. Plan and outline clear and logical steps for a scientific study, and adjust those steps, as necessary, in response to preliminary results
14. Approximately apply the theory of the biology you learn in lecture to the procedures and experiments you do in lab.


Attribution CC-BY-NC-ND.

3.2 Student Attitude towards the Group Environment

*(Adapted from the SAGE instrument by Kouros and Abrami, 2006)*

This questionnaire asks about your attitudes toward small group learning in this classroom. Use your personal experiences from previous classes and group projects to answer these statements.

Please describe your perspective on each statement, on a 5-point scale:
**1= strongly disagree;** 2= disagree; 3= undecided; 4= agree; **5=strongly agree**

1. When I work in a group, I do better quality work.
2. When I work in a group, I end up doing most of the work.
3. When I work with other students, I am able to work at my own pace.
4. When I work in a group, I want to be with my friends.
5. The work takes longer to complete when I work with other students
6. My group members do not respect my opinions.
7. I enjoy the material more when I work with other students
8. My group members help explain things that I do not understand.
9. I become friends with my group members.
10. When I work in a group, I am able to share my ideas.
11. My group members make me feel that I am not as smart as they are.
12. The materials is easier to understand, when I work with other students.
13. My work is better organized, when I am in a group.
14. My group members like to help me learn the material.
15. My group members get a good grade even if they do not do much work.
16. The workload is usually less when I work with other students.
17. I feel I am part of what is going on in the group.
18. One student usually makes the decisions in the group.
19. Our job is not done until everyone has finished the assignment.
20. I find it hard to express my thoughts, when I work in a group.
21. I do not think a group grade is fair.
22. I try to make sure my group members learn the material.
23. My grade depends on how much we all learn.
24. It is difficult to get together outside of class.
25. I learn to work with students who are different from me.
26. My group members do not care about my feelings.
27. I do not like the students I am assigned to work with.
28. I let the other students do most of the work.
29. I get to know my group members well.
30. I feel working in groups, I get the grade I deserve.
31. My group members do not like me.


Attribution CC-BY-NC-ND.

32. I have to work with students who are not as smart as I am.
33. When I work in a group, there are opportunities to express my opinions.
34. When I work with other students, the work is divided equally.
35. My marks improve when I work with other students.
36. Are you still awake? If so, fill in bubble 4 ("agree") to show this.
37. I help my group members with what I am good at.
38. My group members compete to see who does better work.
39. The material is more interesting when I work with other students.
40. When I work in a group, my work habits improve.
41. I like to help my group members learn the material.
42. Some group members forget to do the work.
43. I do not care if my group members get good grades
44. It is important to me that my group gets the work done on time.
45. I am forced to work with students I do not like.
46. I learn more information, when I work with other students.
47. It takes less time to complete the assignment, when I work with others.
48. I also learn when I teach the material to my group members.
49. I become frustrated when my group members do not understand the material.
50. When I work in a group, I get the grade I deserve.
51. Everyone's ideas are needed if we are going to be successful.
52. When I work with other students, we spend too much time talking about other.
53. I prefer to choose the students I work with

3.3 Biology Concept Inventories – please contact the corresponding author for a copy, as not all questions are ours to freely distribute as part of a publication.

3.4 Additional Survey Items – Group Familiarity, Group Interactions, and Opinions

*Pre- and Post-Survey*

- In this lab, having instructor-assigned groups is (choose one: better/worse/no different) than having student-selected groups. Please explain your opinion

*Post-Survey*

1. This semester, how often did you work & interact with your lab group? (1=Labs only, 2=Labs and study time, 3=Labs and lectures; 4= Labs, lectures, and study time)
2. Think back to the start of the course. At the start, how well did you know your group? (1=You had never really met or worked with the other group members before; 2=You knew some of the other group members before, but not all of them; 3=You knew all of them reasonably well (you were friends, and/or you had worked together before.)


Attribution CC-BY-NC-ND.
